# Binding Affinity Estimation From Restrained Umbrella Sampling Simulations

**DOI:** 10.1101/2021.10.28.466324

**Authors:** Vivek Govind Kumar, Shilpi Agrawal, Thallapuranam Krishnaswamy Suresh Kumar, Mahmoud Moradi

**Affiliations:** Department of Chemistry and Biochemistry, University of Arkansas, Fayetteville, AR 72701

## Abstract

The protein-ligand binding affinity quantifies the binding strength between a protein and its ligand. Computer modeling and simulations can be used to estimate the binding affinity or binding free energy using data- or physics-driven methods or a combination thereof. Here, we discuss a purely physics-based sampling approach based on biased molecular dynamics (MD) simulations, which in spirit is similar to the stratification strategy suggested previously by Woo and Roux. The proposed methodology uses umbrella sampling (US) simulations with additional restraints based on collective variables such as the orientation of the ligand. The novel extension of this strategy presented here uses a simplified and more general scheme that can be easily tailored for any system of interest. We estimate the binding affinity of human fibroblast growth factor 1 (hFGF1) to heparin hexasaccharide based on the available crystal structure of the complex as the initial model and four different variations of the proposed method to compare against the experimentally determined binding affinity obtained from isothermal calorimetry (ITC) experiments. Our results indicate that enhanced sampling methods that sample along the ligand-protein distance without restraining other degrees of freedom do not perform as well as those with additional restraint. In particular, restraining the orientation of the ligands plays a crucial role in reaching a reasonable estimate for binding affinity. The general framework presented here provides a flexible scheme for designing practical binding free energy estimation methods.

## INTRODUCTION

Accurate quantification of absolute binding affinities remains a problem of major importance in computational biophysics^1,2,3,4^. In principle, accurate binding free energy calculations should be the cornerstone of any study investigating protein-ligand interactions. However, the high computational costs that typically accompany such calculations necessitate the improvement of the computational methods traditionally used to investigate complex biomolecular interactions^3,4,5^. Experimentally determined binding affinities are commonly used as benchmarks to judge the accuracy of various computational binding affinity estimation methods^5,6,7^. Several experimental techniques can be used to study protein-ligand binding equilibria^5,8^. For instance, isothermal titration calorimetry (ITC) can detect the interaction of binding partners based on changes in solution heat capacity and binding partner concentration^8,9,10^. Other methods such as fluorescence spectroscopy rely on changes in fluorescence intensity upon ligand binding^8,11,12^. Surface plasmon resonance (SPR) can be used to calculate binding affinities based on changes in refractive index that occur when an immobilized binding partner interacts with a free binding partner^8,13,14^. Studies have found that experimental binding affinities can vary depending on the experimental method used^5,6,15^. Therefore, a thorough understanding of the experimental conditions used to generate reference data is essential when comparing computationally determined binding affinities with experimental values.

Several computational methods at varying levels of rigor and complexity have been used to determine binding affinities for biomolecular interactions^3,16-28^. Knowledge-based statistical potentials and force field scoring potentials are typically used to rank docked protein-ligand or protein-protein complexes but can also be used for binding affinity prediction^29,30,31^. A major disadvantage of these methods is that they do not treat the entropic effects rigorously, which effectively decreases the accuracy of such binding affinity predictions^5,32^. This is also the case for methods like Molecular Mechanics/Poisson-Boltzmann-Surface Area (MM-PBSA) and Molecular Mechanics/Generalized Born-Surface Area (MM-GBSA), which combine sampling of conformations from explicit solvent molecular dynamics (MD) simulations with free energy estimation based on implicit continuum solvent models^33,34,35^. Adequate sampling of both ligand conformational dynamics as well as ligand roto-translational movements with respect to the protein is essential for accurately quantifying the entropic reduction arising from the binding event^35,36,37^. MM-PBSA/GBSA methods typically neglect the contribution of these entropic terms to the binding free energy^34,35^.

One of the best-known binding free energy estimation methods is alchemical free energy perturbation (FEP), where scaling of non-bonded interactions enables reversal decoupling of the ligand from its environment in the bound state as well as the unbound state^38,39,40,41^. Most entropic and enthalpic contributors to changes in binding affinity are typically considered during FEP simulations, thus avoiding the approximations used by methods like MM-PBSA/GBSA^5,42^. A disadvantage of FEP is the fact that ligands tend to move away from the binding site during the decoupling process, which results in poorly defined target states of the FEP calculation being used as starting states for the re-coupling process^43^. Using receptor-ligand restraints to resolve this issue^17,40,44,45^ introduces some ambiguity to the way a standard state is defined, with a level of correlation between the size of the simulation cell and the standard state^46^. This can be corrected via the use of appropriate geometrical restraints^16,47,23^.

Unrestrained long timescale MD simulations should theoretically allow for the investigation and accurate quantification of protein-ligand or protein-protein binding events^48,49^. While microsecond-level MD simulations provide a more accurate description of protein conformational dynamics as compared to shorter simulations^50^, efficient sampling of the conformational landscape remains a major issue and requires access to timescales beyond the capabilities of current MD simulations^51,52^. Several methods have been developed to tackle the sampling problem. Markov state models allow the sampling and characterization of native as well as alternative binding states^53,54,55^. Similarly, weighted ensemble (WE) simulations sample the conformational landscape along one or more discretized reaction coordinates based on the assignment of a statistical weight to each simulation^56,57^. More traditionally, umbrella sampling along such reaction coordinates can be used to guide the binding or unbinding of a ligand, after which algorithms like the weighted histogram analysis method (WHAM) can be used to calculate a unidimensional potential of mean force (PMF) which quantifies ligand binding and unbinding along a reaction coordinate^58,59^. Better convergence of the calculated free energy profiles can be achieved by the exchange of conformations between successive umbrella-sampling windows as in the bias-exchange umbrella sampling (BEUS)^60,61,62^. Other methods based on similar principles include umbrella integration^63^, well-tempered metadynamics^64^, adaptive biasing force (ABF) simulations^65^ and variations of these techniques.

Incomplete sampling of important degrees of freedom, such as orientation of the ligand with respect to the protein, remains a major disadvantage of unidimensional PMF-based methods^3,4^. To resolve this problem, Woo and Roux^3^ have devised a method wherein explicitly defined geometrical restraints on the orientation and conformation of the binding partners are used to reduce the conformational entropy of the biomolecular system being studied^3,4^. This results in improved convergence of the PMF calculation^3,4^. The introduction of a restraining potential based on the root-mean-square deviation (RMSD) of the ligand relative to its average bound conformation, reduces the flexibility of the ligand and the number of conformations that need to be sampled^3,4^. This method avoids the need to decouple the ligand from its surrounding environment as required by alchemical FEP^3,4,38-41^. Recent studies have described applications and extensions of the methodology proposed by Woo and Roux^4,66^.

Here, we describe a purely physics-based enhanced sampling method based on biased MD simulations, which is similar in principle to the stratification strategy proposed by Woo and Roux^3,4^. Although we use the US method as our enhanced sampling technique, the methodology is generalizable to other techniques as long as they can be combined with additional restraints. A major difference between our method and that of Woo and Roux^3,4^ is the use of the unidimensional orientation angle of the ligand with respect to the protein as a collective variable for restraining, as opposed to the use of three Euler angles. The formalism has also some other major differences that are discussed in more detail below. We have used use this methodology to calculate the binding affinity for the interaction of human fibroblast growth factor 1 (hFGF1) with heparin hexasaccharide, its glycosaminoglycan (GAG) binding partner. hFGF1 is an important signaling protein that is implicated in physiological processes such as cell proliferation and differentiation, neurogenesis, wound healing, tumor growth and angiogenesis^67,68,69,70,71^. GAGs are linear anionic polysaccharides that interact with positively charged regions of FGF binding partners to regulate their biological activity^70,72-80^. The hFGF1-heparin complex is the most well-known and broadly characterized protein-GAG complex^81,82^. Heparin binding is thought to stabilize hFGF1 and impart protection against proteolytic degradation. In this study, we show that the absolute binding affinity for the hFGF1-heparin interaction calculated using our novel approach, is in good agreement with binding affinity data from ITC experiments.

## THEORETICAL FOUNDATION

Binding affinity is often quantified using the equilibrium dissociation constant (*K*_*d*_), defined as:

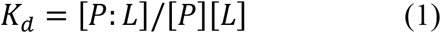

where [*P*], [*L*], and [*P*: *L*] are the concentrations of protein, ligand, and the protein-ligand complex, respectively. Computationally, the absolute binding free energy (Δ*G*°), which is the standard molar free energy of binding, is more convenient to calculate. The dissociation constant and the absolute binding free energy are related via

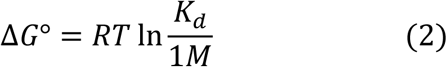

where *R* is the gas constant and *T* is the temperature. Various strategies have been used to estimate Δ*G*°, some of which were briefly discussed above. The methodology proposed here has a significant resemblance to the stratification strategy of Woo and Roux^3,4^. However, the two methods have major differences as will be discussed later.

Absolute binding free energy or Δ*G*° is the free energy change associated with moving the ligand from the bulk to the binding pocket. Within the formalism presented in this work, Δ*G*° is determined from the grid PMF *G*(***x***), where ***x*** is the position of the ligand mass center from the center of the binding pocket, *G*(***x***) is the potential of mean force (PMF) associated with the ligand position ***x***. In practice, we need to bin the 3D space and define the PMF at every bin or grid point as:

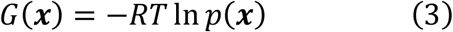

where *p*(***x***) is the probability of finding the ligand at bin ***x***.

We define Δ*G*(***x***) = *G*(***x***) − *G*(**0**), where ***x*** = **0** (i.e., the center of the binding pocket) is defined as the grid point associated with the lowest grid PMF. One can show:

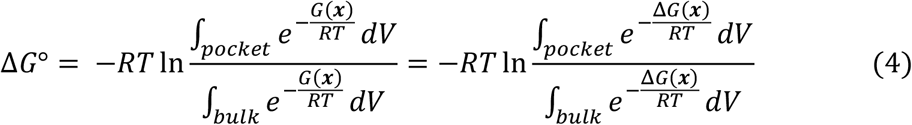

in which the pocket refers to all ***x*** where the ligand is considered bound (i.e., the binding pocket) and bulk refers to all ***x*** where the ligand is not interacting with the protein. Since Δ*G*(***x***) is the same everywhere in the bulk, we can simplify Relation (4) as follows:

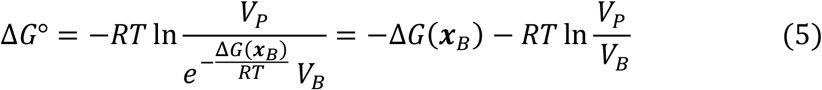

where *V*_*B*_ is the bulk volume per protein associated with the standard concentration (i.e., 1 *M*), ***x***_*B*_ is any grid point in the bulk, and *V*_*P*_ is the binding pocket volume defined as:

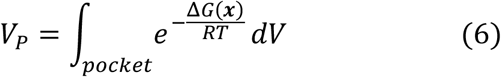

Defining Δ*G*_*V*_ as the contribution of the difference between the volume of the binding pocket and the bulk to the binding free energy:

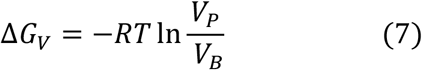

Combining (5) and (7), we have:

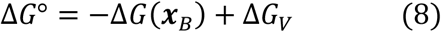

We can find the bulk volume (*V*_*B*_) associated with the standard concentration for a single protein approximately as:

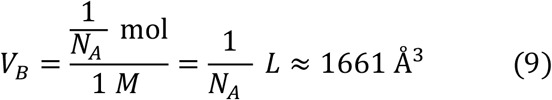

where ***N***_*A*_ is the Avogadro’s constant. We can now rewrite Δ*G*_*V*_ as:

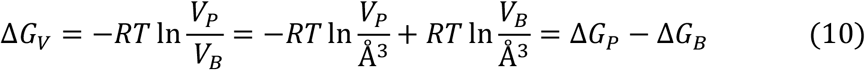

in which Δ*G*_*B*_ is the bulk volume contribution and Δ*G*_*P*_ is the binding pocket contribution:

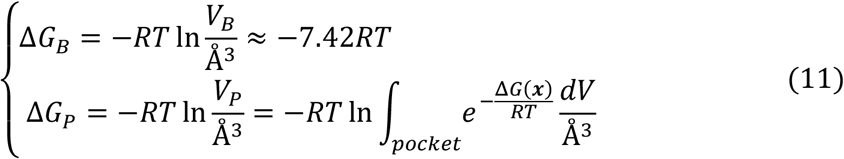

Determining both Δ*G*(***x***_*B*_) and Δ*G*_*P*_ requires finding the grid PMF Δ*G*(***x***). Δ*G*(***x***_*B*_) is the PMF difference between the binding pocket center and the bulk and Δ*G*_*P*_ also requires an estimate for Δ*G*(***x***) within the binding pocket. We therefore do not need to find Δ*G*(***x***) for all ***x*** if we have a good estimate for Δ*G*(***x***) within the binding pocket and in the bulk. Ideally, Δ*G*(***x***) for these points can be determined by pulling the ligand out of the binding pocket towards the bulk and using an enhanced sampling technique such as US to sample the space of a collective variable such as *d*, i.e., the distance between the mass centers of the ligand and protein. Δ*G*(***x***) can be estimated for all sampled grid points ***x*** using this distance-based US simulation. Note that the collective variable used for biasing would be *d*, while the collective variable used for the PMF calculations would be the 3D position vector of the mass center of ligand with respect to protein’s binding pocket center. One may estimate the grid PMF from the distance-based US simulations using a non-parametric reweighting algorithm as discussed in the Methods section. Δ*G*(***x***) can also be used to estimate Δ*G*_*P*_ as defined in Relation (11). There is often no need to strictly define the binding pocket since only low Δ*G*(***x***) values have nonnegligible contribution to *V*_*P*_ and thus even if we include all sampled grid points, only those close to the binding pocket center have nonnegligible contributions.

A practical issue with determining Δ*G*(***x***_*B*_) is the convergence. The key obstacles for the sampling that slow down the convergence are the orientation of the ligand, and the conformational changes of the ligand and protein. Using an approach similar in spirit to the previously proposed stratification strategy^3,4,35^, we can circumvent extensive sampling of these degrees of freedom. Let us first focus on the orientation of the ligand (Ω). We can restrain Ω during the distance-based US simulations using a biasing potential 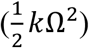 and later correct the free energy difference based on the PMF associated with the Ω, which is different in the bulk (*F*(***x***_B_, Ω)) and in the binding pocket (*F*(**0**, Ω)). More generally, for any grid point ***x***, we may determine Δ*G*(***x***) based on the PMF associated with the Ω at ***x*** (*F*(***x***, Ω)) and **0** (*F*(**0**, Ω)):

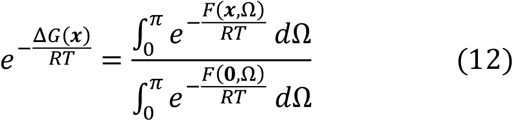

Note that *F*(***x***, Ω) is the PMF associated with ***x*** and Ω, defined such that:

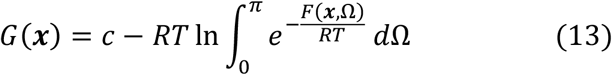

where *c* is an arbitrary constant. We therefore have:

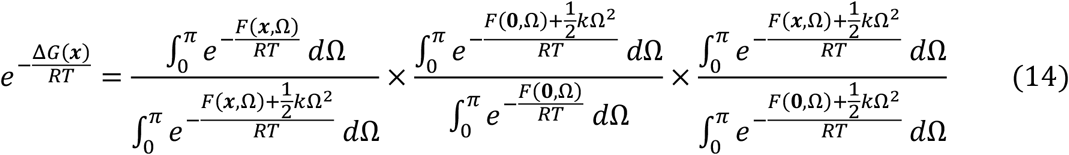

We now define *G*_Ω_(***x***) as the grid PMF of the restrained system (by Ω):

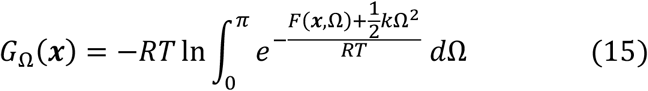

We also define *U*_Ω_(***x***) as the average biasing potential at grid point ***x***:

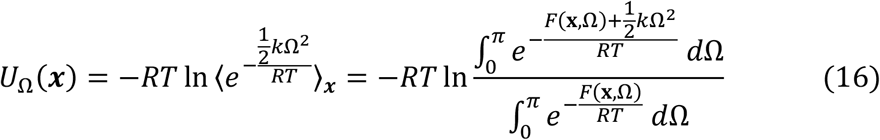

Now we have from Relations (14), (15), and (16):

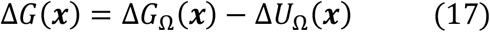

where the free energy of grid point ***x*** from the center **0** (Δ*G*(***x***)) is calculated based on its equivalent free energy (Δ*G*_Ω_(***x***)) in a system biased by a harmonic restraint on Ω and a correction term Δ*U*_Ω_(***x***). For ***x*** = ***x***_*B*_:

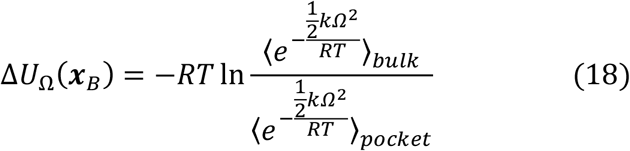

To determine the above ensemble averages, we need to determine the PMF along *Ω* for the bound and unbound ligand and calculate the ensemble averages analytically using Relation (16). Δ*G*_Ω_(***x***_*B*_) can be determined from PMF calculations, where the distance between the protein and ligand is varied and the orientation of the ligand is restrained (distance-based BEUS with restrained orientation). We note that:

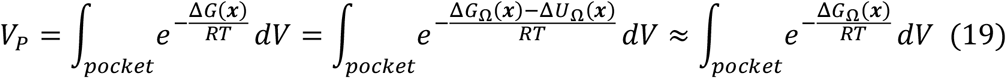

where we assume Δ*U*_Ω_(***x***) is negligible for ***x*** within the binding pocket. In other words, 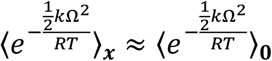 for ***x*** close to **0**.

In brief, if we choose to restrain the orientation, our absolute binding free energy estimate includes the following terms (using Relations (8) and (17)):

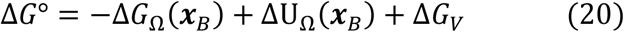

*F*(***x***_*B*_, Ω) can be calculated numerically from orientation angle distribution of a free ligand: F(***x***_*B*_, Ω) = − *RT* ln *p*(Ω), where *p*(Ω) is determined from the distribution of Euler angles 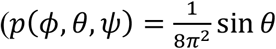, where 0 ≤ *ϕ, Ψ* ≤ 2*π* and 0 ≤ *θ* ≤ *π*) given that:

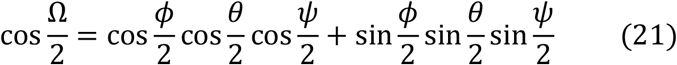

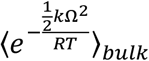 can then be calculated using numerically calculated *F*(***x***_*B*_, Ω) and *k* as used in the simulations using Relation (16). *F*(**0**, Ω) can be determined approximately using orientation-based US simulations of bound ligand. F(**0**, Ω) can then be used to estimate 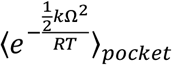 using Relation (16).

The above strategy can be extended to other degrees of freedom for which unbiased sampling may hinder the convergence. Most notably, the internal conformational changes of the ligand and that of the protein may also play a crucial role in slowing down the convergence. In the following, we show how one can restrain not only the orientation of the ligand but also the root-mean-square-deviation (RMSD) of the ligand (denoted here by *r*) in distance-based US simulations (along *d*) to speed up convergence. In this case, the grid PMF difference Δ*G*(***x***) is calculated based on Δ*G*_Ω,*r*_(***x***), the grid PMF of a system whose Ω and *r* are both restrained:

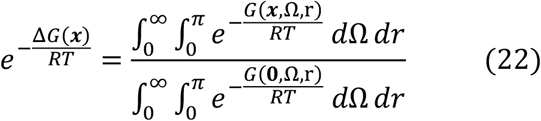

Using a similar strategy as in Relation (14), we have:

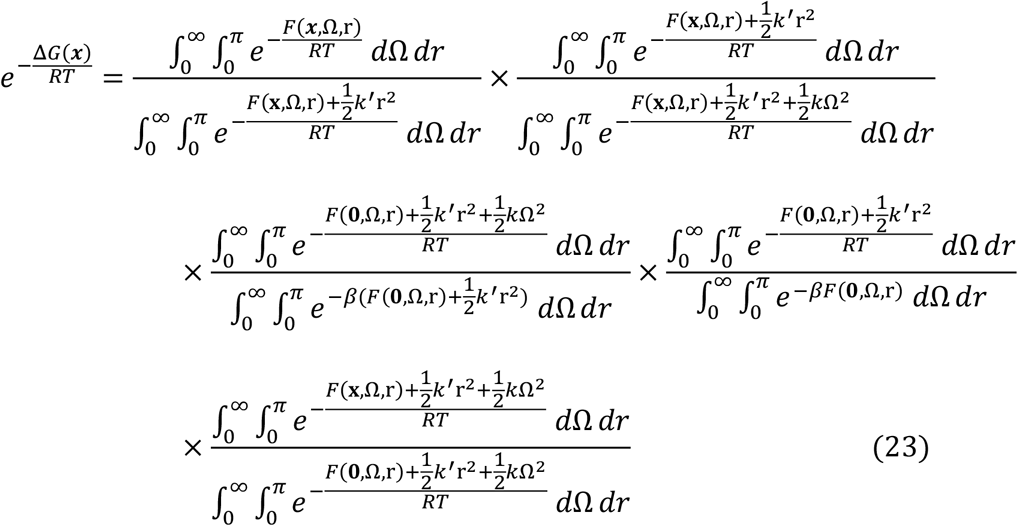

which results in:

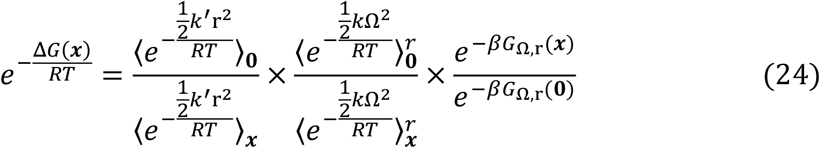

Here we have defined *G*_Ω,r_(***x***) as:

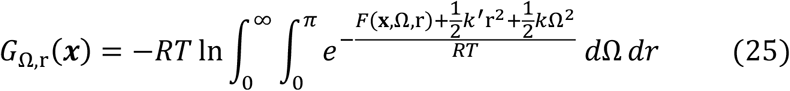

We also define *U*_*r*_(***x***) similar to *U*_Ω_(***x***) in Relation (15) except for using *r* instead of Ω. 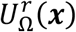 is also defined similar to *U*_Ω_(***x***) except for the additional restraint on *r*:

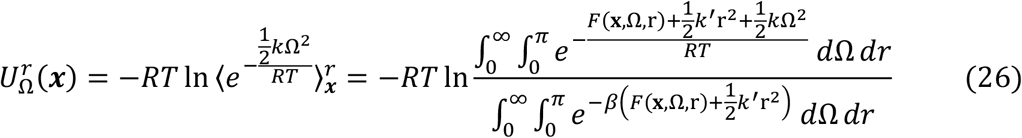

Finally, we have:

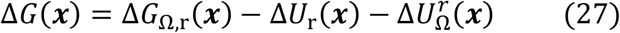

In brief, if we choose to restrain both the orientation and RMSD, our absolute binding free energy estimate includes the following terms:

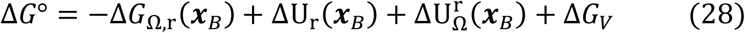

Here we are using an approximation similar to that in Relation (19):

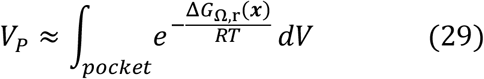

Using Relations (20) and (28), we can generalize the stratification strategy to include three restraints on arbitrary collective variables *α, β*, and *γ*:

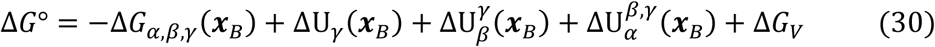

where:

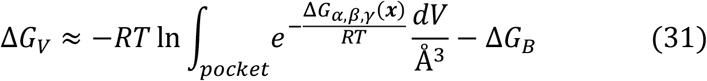

## METHODS

### Isothermal titration calorimetry of hFGF-1 with heparin hexasaccharide

Isothermal titration calorimetry (ITC) data was obtained using MicroCal iTC 200 (Malvern Inc.). The change in heat during the biomolecular interaction was measured by titrating the heparin (loaded in the syringe) to the hFGF1 solution in the calorimetric cell. Both the protein and the heparin samples were made in the buffer containing 10 mM phosphate buffer with 100 mM NaCl at pH 7.2 and were degassed prior to loading. The protein to heparin ratio was maintained at 1:10 with the protein concentration being 100 μM and the heparin concentration being 1mM. A total of 30 injections were conducted with a constant temperature of 25 °C and stirring speed of 300 rpm. One set of sites binding model was used for the ITC binding curve^85^. The standard binding free energy ΔG° was determined from dissociation constant via Relation (2) at *T* = 25°C.

### All-atom MD simulations

Our simulations were based on the x-ray crystal structure of the dimeric complex with a heparin hexasaccharide (PDB:2AXM, resolution: 3.0 angstroms)^86^. One of the hFGF1 protomers was removed leaving one protein and one ligand in the model of the holo protein. The model for the apo protein was based on the x-ray crystal structure of unbound monomeric hFGF1 (PDB: 1RG8, resolution: 1.1 angstroms)^87^. The simulations (residues 12-137 in the PDB file correspond to residues 26-151 in the experimental sequence) and experiments were performed using a truncated version of hFGF1 (residues 13-154) which did not contain the unstructured 12 amino acid N-terminal segment. The heparin hexasaccharide consists of N, O6 disulfo-glucosamine and 2-O-sulfo-alpha-L-idopyranuronic acid repeats^86^. The models were solvated in a box of TIP3P waters and 0.15 M NaCl. MD simulations were performed using the NAMD 2.13^88^ simulation package with the CHARMM36m all-atom additive force field^89^ Initially, we energy-minimized the systems for 10,000 steps using the conjugate gradient algorithm^90^. Subsequently, we relaxed the systems using restrained MD simulations in a stepwise manner (for a total of ∼1 ns) using the standard CHARMM-GUI protocol^91,92^. The initial relaxations were performed in an NVT ensemble while the production runs were performed in an NPT ensemble. Simulations were carried out using a 2-fs time step at 300 K using a Langevin integrator with a damping coefficient of γ = 0.5 ps^−1^. The pressure was maintained at 1 atm using the Nosé−Hoover Langevin piston method^90,93^. The smoothed cutoff distance for non-bonded interactions was set to 10−12 Å and long-range electrostatic interactions were computed with the particle mesh Ewald (PME) method^94^. The initial runs lasted 15 nanoseconds, followed by the productions run on the supercomputer Anton 2 (Pittsburgh Supercomputing Center) for 4.8 μs, with a timestep of 2.5 fs. These equilibrium simulations have previously been described in a related study^95^. We used these equilibrium simulations to construct the PMF in terms of the RMSD of the protein (*r*_*P*_) both for the apo^95^ and holo proteins (for bulk and binding pocket, respectively). We also used the holo protein simulations^95^ to construct the PMF in terms of the RMSD of the ligand (*r*_*L*_) in the binding pocket.

### MD simulations of free heparin hexasaccharide

The heparin hexasaccharide^86^ was simulated in a rectangular water box without the protein. The system was set up as described previously. The final conformation after relaxation was then used as the starting conformation for 10 production runs for 40 ns each. The total simulation time was around 400 ns. We used these unbiased simulations instead of US simulations to construct the PMF of free heparin in the bulk in terms of ligand RMSD (*r*_*L*_).

### Steered Molecular Dynamics (SMD) simulations

The final conformation of the hFGF1-heparin equilibrium simulation^95^ was used to generate starting conformations for the non-equilibrium pulling simulations. Two collective variables^96^ were used for SMD simulations^97^: (1) distance between the heavy-atom center of mass of heparin and that of the protein (*d*) and (2) the orientation angle of heparin with respect to the protein (Ω). Two independent sets of simulations were performed. The distance-based SMD simulation was run for 9.5 ns, while the orientation based SMD simulation was run for 8 ns. The distance-based SMD simulation was used to pull the heparin away from the protein by approximately 30 Å (10→40 Å) with a force constant of 100 kcal/(mol. Å^2^). The orientation angle was also restrained in these simulations with a force constant of 0.5 kcal/(mol.*degree*^2^) to stay close to its initial orientation in the bound state. The orientation-based SMD simulation was used to rotate the bound heparin locally with respect to the protein (0°→73°) with a force constant of 100 kcal/(mol.*degree*^2^).

### Bias Exchange Umbrella Sampling (BEUS) simulations

Bias exchange umbrella sampling^62,98,99^ (BEUS), which is a variation of the US simulation method, was performed to estimate grid PMF. Four independent sets of distance (*d*) based BEUS simulations were performed, with no restraints, restraint on Ω, restraint on *r*_*L*_ and *r*_*P*_, and restraints on Ω, *r*_*L*_, and *r*_*P*_. Two sets of BEUS simulations were also performed using the Ω collective variable, one with and one without a restraint on *r*_*L*_ and *r*_*P*_. Selected SMD conformations were assigned to individual BEUS windows with equal spacing in each one of these BEUS simulations. The distance-based BEUS simulation ran for 10 ns with 31 replicas/windows and the orientation-based simulation ran for 10 ns with 30 replicas/windows. The force constant used for ligand-protein distance (*d*) in distance-based BEUS was 2 kcal/(mol.Å^2^) while the orientation was restrained as in SMD simulations using a force constant of 0.5 kcal/(mol. *degree*^2^). For orientation-based BEUS simulations, the force constant for the ligand orientation angle (as in SMD simulations) was set to 0.5 kcal/(mol. *degree*^2^). The force constant used for *r*_*L*_ and *r*_*P*_ was 1 kcal/(mol.Å^2^).

### Free energy calculations using non-parametric reweighting

Once the BEUS simulations described above were converged, a non-parametric reweighting method^98,100^, which is somewhat similar to the multi-state Bennett acceptance ratio method^101^, was used to construct PMF. In this method^98^, each sampled configuration will be assigned a weight, which can be used to construct the PMF in terms of a desired collective variable. Suppose that a system is biased (for instance, within a BEUS scheme) using N different biasing potentials *U*_*i*_(***r***), where *i* = 1, …, ***N***, and ***r*** represents all atomic coordinates. Typically, *U*_*i*_(***r***) is a harmonic potential defined in terms of a collective variable with varying centers for different *i*. Assuming an equal number of sampled configurations from each of the ***N*** generated trajectories, we can combine them in a single set of samples {***r***_*k*_} (irrespective of which bias was used to generate each sample ***r***_*k*_) and determine the weight of each sample as:

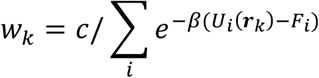

where *c* is the normalization constant such that ∑_*k*_ *w*_*k*_ = 1 and both {*w*_*k*_} and {*F*_*i*_} are determined iteratively using the above equation and the following:

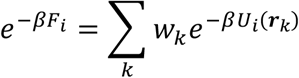

Converged *w*_*k*_ values can be used to construct any ensemble averages including any PMF (e.g., *G*(***ζ***)) not only in terms of the collective variable used for biasing but also any other collective variables that are sufficiently sampled. One may use a weighted histogram method to construct the PMF as follows:

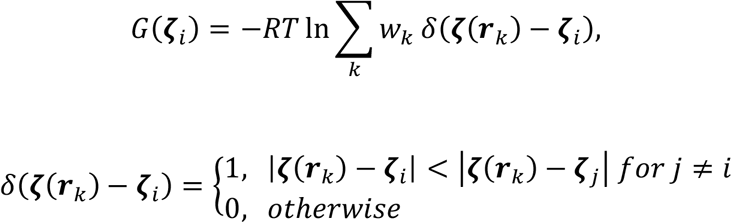

## RESULTS AND DISCUSSION

We have calculated the absolute binding free energy for the interaction of hFGF1 with heparin hexasaccharide using four variations of the stratification scheme described above, based on a combination of SMD and BEUS simulations. The details of the methodology are discussed in the Methods section. Four different methods are used with varying effectiveness in estimating the absolute binding free energy. These methods include (1) the traditional distance-based BEUS simulations that do not employ any additional restraining, (2) distance-based BEUS simulations employing a restraint on the orientation of the ligand (Ω) defined based on the orientation quaternion, (3) distance-based BEUS simulations employing a restraint on the RMSD of both ligand and protein (r_L_, *r*_*P*_), (4) distance-based BEUS simulations employing a restraint on the RMSD of both ligand and protein as well as the orientation of the ligand (Ω, r_L_, *r*_*P*_). In each case, appropriate correction terms are calculated as discussed in the Theoretical Foundation section above and shown in Table 1.

**Table 1:** Summary of free energy calculation results.

| | No restraints | $\Omega$ restraint | $r_L, r_P$ restraint | $\Omega, r_L, r_P$ restraint |
| --- | --- | --- | --- | --- |
| Grid PMF difference<br>(kcal/mol) | $\Delta G(\mathbf{x}_B) =$<br>$-19.7 \pm 1.1^*$ | $\Delta G_\Omega(\mathbf{x}_B) =$<br>$-13.2 \pm 0.3$ | $\Delta G_{r_L, r_P}(\mathbf{x}_B) =$<br>$-17.7 \pm 1.0$ | $\Delta G_{\Omega, r_L, r_P}(\mathbf{x}_B) =$<br>$-17.0 \pm 0.5$ |
| Orientation correction<br>(kcal/mol) | N/A | $\Delta U_\Omega(\mathbf{x}_B) =$<br>$4.4 \pm 0.3$ | N/A | $\Delta U_\Omega^{r_L, r_P}(\mathbf{x}_B) =$<br>$4.4 \pm 0.3$ |
| Ligand RMSD correction<br>(kcal/mol) | N/A | N/A | $\Delta U_{r_L}(\mathbf{x}_B) =$<br>$0.5 \pm 0.1$ | $\Delta U_{r_L}(\mathbf{x}_B) =$<br>$0.5 \pm 0.1$ |
| Protein RMSD correction<br>(kcal/mol) | N/A | N/A | $\Delta U_{r_P}^{r_L}(\mathbf{x}_B) =$<br>$0.8 \pm 0.1$ | $\Delta U_{r_P}^{r_L}(\mathbf{x}_B) =$<br>$0.8 \pm 0.1$ |
| $\Delta G_V$ (kcal/mol) | $3.7 \pm 0.2$ | $2.5 \pm 0.2$ | $2.3 \pm 0.2$ | $2.7 \pm 0.2$ |
| $\Delta G^\circ$ (kcal/mol) | $-16.0 \pm 1.2$ | $-6.3 \pm 0.5$ | $-14.1 \pm 1.0$ | $-8.5 \pm 0.7$ |
| $K_d$ ($\mu M$)<sup>**</sup> | $O(10^{-6})$ | 25 | $O(10^{-5})$ | 0.6 |
| $K_d$ range ($\mu M$)<sup>***</sup> | $10^{-7} - 10^{-5}$ | 11 – 58 | $10^{-6} - 10^{-4}$ | 0.2 – 2.0 |

\* All error estimates are based on one standard deviation (s.d.). \*\* $K_d$ values are determined directly from mean $\Delta G^\circ$ values using Relation (2). \*\*\* $K_d$ range is determined from the lower and upper limits of $\Delta G^\circ$ values (mean ± s.d.) using Relation (2).

The most successful method is expected to be the one employing restraints on Ω, r_L_, *r*_*P*_. The largest contributor to the free energy is the difference between the grid PMF associated with the heparin hexasaccharide at a grid point at the center of the binding pocket and at any grid point in the bulk, which is -17.0 ± 0.5 kcal/mol (see Figure 1A and Table 1). We denote the PMF of the ligand at a given position ***x*** (with respect to the center of the heparin binding pocket) as the grid PMF, since the PMF is estimated at different grid points in this approach.

**Figure 1.**
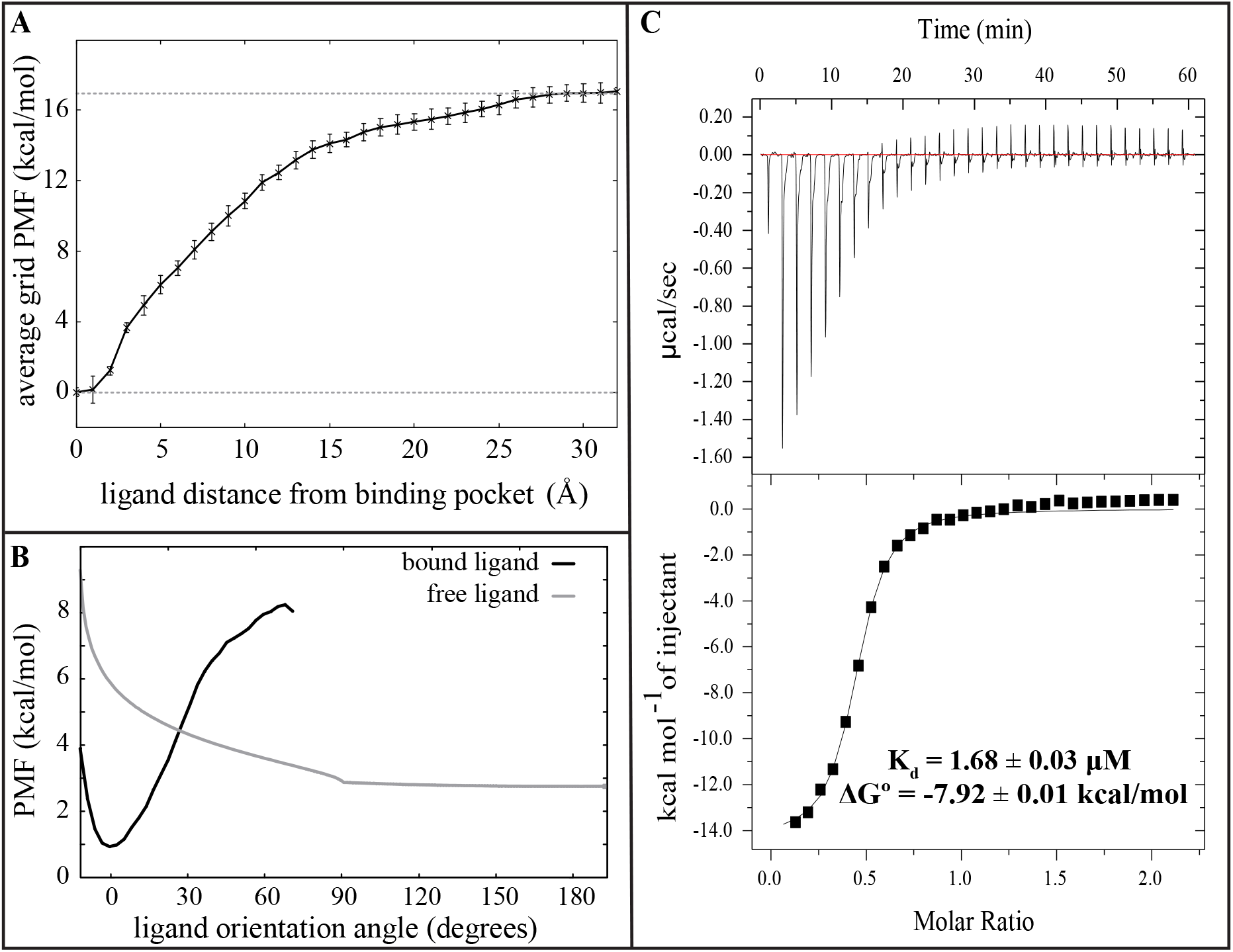
Computational (A-B) and experimental (C) heparin-hFGF1 binding free energy measurements. (A) Average grid PMF in terms of |***x***|,where ***x*** is the 3D position vector of the ligand with respect to the center of binding pocket determined from distance-based BEUS simulations with Ω, *r*_*L*_, *r*_*P*_ restraints. The x axis represents |***x***| and the y axis represents 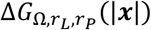, which is an average over all 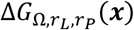 with the same |***x***|, i.e., the ligand distance from the center of binding pocket. The error bar represents the standard deviation obtained from all values of 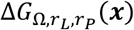 at various grid points ***x*** with the same |***x***|. The dashed line represents the value associated with 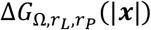 at |***x***| = 30 Å. (B) The PMF associated with the ligand orientation angle (Ω) for the bound heparin (i.e., ***x*** ≈ **0**, ligand in the binding pocket) and free heparin (i.e., ***x*** ≈ ***x***_*B*_, ligand in the bulk). (C) Isothermogram representing the titration of hFGF1 with heparin hexasaccharide. The inset is the experimentally estimated dissociation constant and its associated binding free energy.

The PMF calculations above are based on the BEUS simulations along the protein-ligand distance; however, the orientation and RMSD of the ligand and the RMSD of the protein are restrained to speed up convergence. To account for the orientation bias, a correction term needs to be applied, which is calculated from the PMF associated with the ligand orientation angle at the bulk and binding pocket (Figure 1B). The orientation bias is estimated to be 4.4 ± 0.3 kcal/mol (Table 1). Similarly, a correction term is calculated based on the PMF of the ligand RMSD and that of the protein (Figure 2). These correction terms are estimated to be 0.5 ± 0.1 and 0.8 ± 0.1 kcal/mol, for the ligand and protein, respectively.

**Figure 2.**
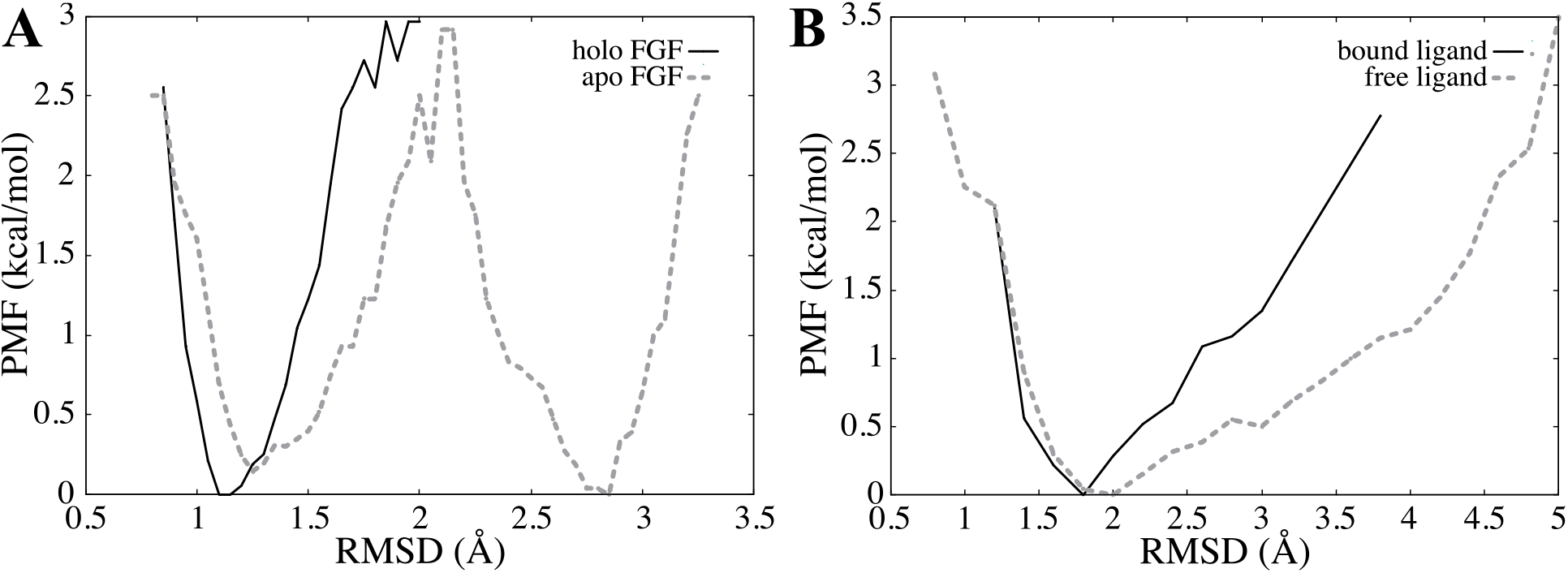
PMF in terms of internal conformational fluctuations of the protein and ligand. (A) PMF associated with the internal RMSD of heparin-bound (solid line) and apo (dashed line) hFGF1, obtained from the equilibrium simulations. (D) PMF associated with the internal RMSD of FGF1-bound (solid line) and free (dashed line) heparin hexasaccharide, obtained from equilibrium simulations.

Finally, another term is needed to account for the difference in the volume accessible to the ligand in the binding pocket and in the bulk (volume contribution). Figure 3 shows that Δ*G*_*P*_ (or *V*_*P*_) for the distance-based BEUS simulations with no restraint as determined from 20 lowest free energy grid points is almost equal to that obtained from all visited grid points inside or outside the binding pocket. For the distance-based BEUS simulations with Ω, r_L_, *r*_*P*_ restraints, this term is estimated to be 2.7 ± 0.2 kcal/mol, which results in an absolute binding free energy of -8.5 ± 0.7 kcal/mol. Based on our error analysis, K_d_ values calculated from the absolute binding free energy were found to be in the micromolar range with an average value of 0.6 μM (using the mean Δ*G*° estimate) and ranging from 0.2 to 2.0 μM (based on the lower and upper bounds of free energy estimates). These are in very good agreement with the K_d_ value obtained from ITC experiments that is 1.68 μM (Figure 1C). The free energy calculated from the experimental K_d_ (−7.91 kcal/mol) is also in good agreement with the computationally calculated binding free energy (Figure 1C and Table 1).

**Figure 3.**
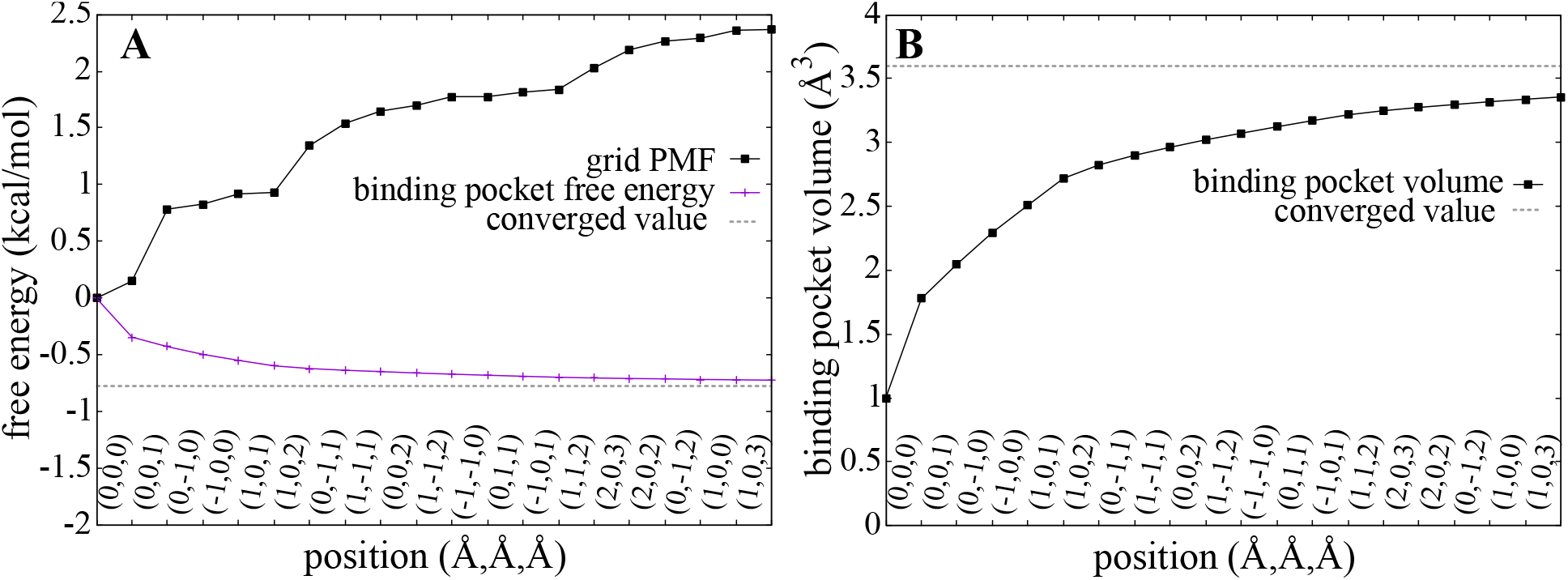
Estimating binding pocket volume (*V*_*P*_) and its contribution to absolute binding free energy (Δ*G*_*P*_). (A) Grid PMF (Δ*G*(***x***)) associated with grid points with the 20 lowest PMF values (black) along with estimated Δ*G*_*P*_ based on the first 20 grid points (shown in an accumulative manner in magenta). The distance-based BEUS simulations with no restraints are used for these calculations. Dashed line shows the estimated Δ*G*_*P*_ based on all visited grid points inside or outside the binding pocket. The x axis shows the position vector of these 20 grid points. (B) Binding pocket volume (*V*_*P*_) calculated from 20 lowest grid PMF values (similar to A). Dashed line shows the *V*_*P*_ estimated from all visited grid points inside or outside the binding pocket. See the Methods section for more details.

The quantitative agreement between the computational and experimental binding affinity estimates is a great indicator of the accuracy of our absolute binding free energy calculation method. However, if proper restraining is not used as in the distance-based BEUS simulations with no restraints or only RMSD restraints, the binding affinity estimates would be off by several orders of magnitude. The simulations that only restrain the orientation of the ligand are interestingly quite successful as well, being off only by one order of magnitude in terms of binding affinity, which is generally considered a good estimate. This provides some evidence that the orientation of the ligand is perhaps the degree of freedom with the most significant contribution to the absolute binding free energy besides the ligand-protein distance. We note that the average grid PMF profiles along the ligand-protein distance for the four different methods used here (as shown in Figure 4), confirm the differential behavior of these methods; however, it is important to note that the correction terms should ideally eliminate these differences. This is seen to some extent when comparing the two methods involving orientation restraints that happen to estimate binding affinities that are reasonably close to the experimentally determined value.

**Figure 4.**
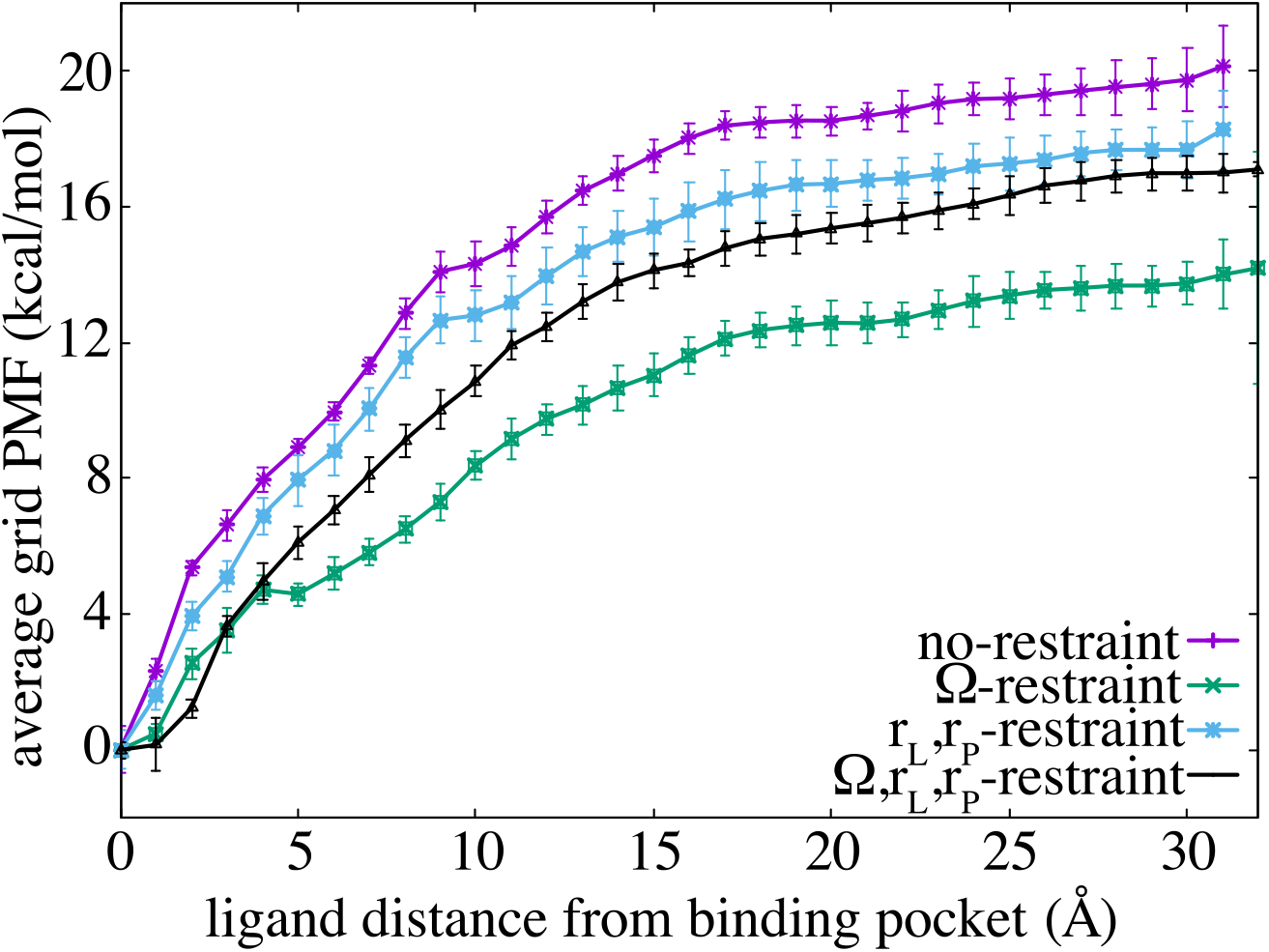
Average grid PMF based on four alternative distance-based BEUS simulations. Average grid PMF in terms of |***x***|,where ***x*** is the 3D position vector of the ligand with respect to the center of binding pocket. The x axis represents |***x***| and the y axis represents Δ*F*(|***x***|), which is an average over all Δ*F*(***x***) with the same |***x***|, i.e., the ligand distance from the center of binding pocket. The error bar represents the standard deviation obtained from all values of Δ*F*(***x***) at various grid points ***x*** with the same |***x***|. The dashed line represents the value associated with Δ*F*(|***x***|) at |***x***| = 30 Å. The inset summarizes different free energy terms involved in the calculation of the absolute binding free energy and the dissociation constant.

Recent computational studies have used the MM-GBSA method to calculate the binding free energy of the hFGF1-heparin interaction, with values ranging from -84.9 kcal/mol to -106.1 kcal/mol^102^. The results obtained from the MM-GBSA approach are very different from our own results, which is to be expected given that MM-GBSA ignores various contributors to the free energy^33,34,35^. Studies have shown that the binding affinity and free energy results derived from computational methods can be compared to experimental binding affinities obtained from ITC experiments^9,10^. However, for a reliable computational free energy estimate, employing purely physics-based free energy calculation methods such as those employed here has proven to be difficult. Here we showed that using a careful strategy that considers all relevant free energy terms and ensures the use of powerful enhanced sampling techniques, could result in good quantitative agreements between the computational and experimental binding affinity estimates.

The formalism presented in this work has significant similarities to the method previously proposed by Woo and Roux^3^, and later implemented^4,66^. However, there are major differences that make the current method more practical. The grid PMF and its various estimates provide a simple conceptual framework to understand how restraining can be accounted for with appropriate correction terms. The average grid PMF in terms of the ligand-protein distance provides an alternative to the PMF in terms of *d* as is often constructed. Relation (30) is a general scheme that can be easily adapted to any number of restraints. The orientation angle of the ligand with respect to the protein as determined using the orientation quaternion formalism, provides a simple way of determining the absolute binding free energy with a feasible computational cost. Among the four different sets of restraints, the two involving orientation restraints predict binding free energies similar to that determined experimentally.

## ACKNOWLEDGEMENTS

This research is supported by National Science Foundation grant CHE 1945465 and OAC 1940188 and the Arkansas Biosciences Institute. Anton 2 computer time was provided by the Pittsburgh Supercomputing Center (PSC) through Grant R01GM116961 from the National Institutes of Health. The Anton 2 machine at PSC was generously made available by D.E. Shaw Research. This research is also part of the Blue Waters sustained-petascale computing project, which is supported by the National Science Foundation (awards OCI-0725070 and ACI-1238993) and the state of Illinois. This work also used the Extreme Science and Engineering Discovery Environment (allocation MCB150129), which is supported by National Science Foundation grant number ACI-1548562. This research is also supported by the Arkansas High Performance Computing Center, which is funded through multiple National Science Foundation grants and the Arkansas Economic Development Commission. This work is also supported by the Department of Energy (DE-FG02-01ER15161), the National Institutes of Health/National Cancer Institute (NIH/NCI) (1 RO1 CA 172631) and the NIH through the COBRE program (P30 GM103450).

## AUTHOR CONTRIBUTIONS

M.M. and T.K.S.K. designed the research. V.G.K performed the simulations and analyzed the simulation data. S.A. performed the experiments and analyzed the experimental data. V.G.K, M.M., T.K.S.K., S.A. wrote the manuscript.

## AVAILABILITY OF DATA

All data will be shared upon request to corresponding author.

